# Leveraging GPT-4 for Identifying Cancer Phenotypes in Electronic Health Records: A Performance Comparison between GPT-4, GPT-3.5-turbo, Flan-T5 and spaCy’s Rule-based & Machine Learning-based methods

**DOI:** 10.1101/2023.09.27.559788

**Authors:** Kriti Bhattarai, Inez Y. Oh, Jonathan Moran Sierra, Jonathan Tang, Philip R.O. Payne, Zachary B. Abrams, Albert M. Lai

## Abstract

**Objective:** Accurately identifying clinical phenotypes from Electronic Health Records (EHRs) provides additional insights into patients’ health, especially when such information is unavailable in structured data. This study evaluates the application of OpenAI’s Generative Pre-trained Transformer (GPT)-4 model to identify clinical phenotypes from EHR text in non-small cell lung cancer (NSCLC) patients. The goal was to identify disease stages, treatments and progression utilizing GPT-4, and compare its performance against GPT-3.5-turbo, Flan-T5-xl, Flan-T5-xxl, and two rule-based and machine learning-based methods, namely, scispaCy and medspaCy.

**Materials and Methods:** Phenotypes such as initial cancer stage, initial treatment, evidence of cancer recurrence, and affected organs during recurrence were identified from 13,646 records for 63 NSCLC patients from Washington University in St. Louis, Missouri. The performance of the GPT-4 model is evaluated against GPT-3.5-turbo, Flan-T5-xxl, Flan-T5-xl, medspaCy and scispaCy by comparing precision, recall, and micro-F1 scores.

**Results:** GPT-4 achieved higher F1 score, precision, and recall compared to Flan-T5-xl, Flan-T5-xxl, medspaCy and scispaCy’s models. GPT-3.5-turbo performed similarly to that of GPT-4. GPT and Flan-T5 models were not constrained by explicit rule requirements for contextual pattern recognition. SpaCy models relied on predefined patterns, leading to their suboptimal performance.

**Discussion and Conclusion:** GPT-4 improves clinical phenotype identification due to its robust pre-training and remarkable pattern recognition capability on the embedded tokens. It demonstrates data-driven effectiveness even with limited context in the input. While rule-based models remain useful for some tasks, GPT models offer improved contextual understanding of the text, and robust clinical phenotype extraction.

## BACKGROUND AND SIGNIFICANCE

### Introduction

Extracting clinical phenotypes from unstructured Electronic Health Records (EHRs) is a critical task in natural language processing (NLP). Accurately identifying relevant phenotypes from unstructured text utilizing NLP techniques provides additional insights into patients’ health, especially when such information is unavailable in structured data. NLP extraction techniques facilitate this process by mapping unstructured text to a structured representation, making it easier to evaluate patients’ disease progression, treatment modalities, and treatment effectiveness. This is particularly evident when analyzing data from non-small cell lung cancer patients, where unstructured text is abundant. Accurately identifying disease stage, treatments and progression from cancer text will contribute to continued research efforts aimed at improving treatment strategies for non-small lung cancer patients, assessing disease progression, and improving lung cancer-related outcomes.

### Background

Clinical phenotype extraction is an ongoing research area where the type of extraction tasks and target phenotypes vary across different clinical domains. Rule-based, machine learning-based, and deep-learning models have been applied to phenotype extraction.^1-7^ While rule-based models extract phenotypes based on pre-defined patterns, most machine learning and deep-learning approaches are trained on sentences or documents labeled with the relevant phenotypes and the model subsequently classifies texts into these phenotypes.^5,8^ SpaCy models, including MedspaCy^7^ and scispaCy^9^ are two recent and frequently used hybrid frameworks that utilize statistical and machine-learning named entity recognition methods in conjunction with rule-based NLP to identify clinical phenotypes. There are studies that have utilized medspaCy and scispaCy to identify specific sections within EHR text for NER, extract phenotypes from relation extraction documents, and generate text embeddings.^10-14^

Although extracting clinical phenotypes is essential, several gaps remain in the literature. There is no effective model for direct extraction, as most of these models require additional training and fine-tuning.^15,16^ Moreover, current methods often lack robustness, leading to suboptimal performance.^15-19^ In addition, limited availability of labeled, publicly accessible cancer EHR text leaves an important domain underexplored for NLP.

Pre-trained transformer-based language models have recently been studied for tasks such as question answering, text generation, and machine translation.^20,21^ Despite the success of transformer-based language model in such tasks, their application in the context of clinical phenotype extraction remain underexplored, opening numerous avenues of research. Recent research has demonstrated the use of large language models (LLMs) for entity extraction.^22-25^ However, it is essential to investigate these recent transformer-based methods in specific clinical domains and compare their performance to previously recognized machine learning and rule-based models to generate insights into their potential benefits for clinical phenotype extraction.

## OBJECTIVES

The aim of this study was to investigate the most recent transformer-based language models as they remain underexplored for cancer phenotype extraction from real-world EHR text. We evaluated the application of OpenAI’s Generative Pre-Trained Transformer (GPT)-4 model^25^ for clinical phenotype extraction in an EHR retrospective study focusing on non-small cell lung cancer patients as a specific case study. We used GPT-4 to identify individual words or tokens in a data sequence as distinct phenotypes. Specifically, we measure the prevalence of specific lung cancer phenotypes, including cancer stage, treatment modalities, cancer recurrence instance, and organs affected by cancer recurrence. These phenotypes are important for informing treatment decisions and assessing disease progression in non-small cell lung cancer patients.

We built the model framework using a clinical text dataset from Washington University in St. Louis, Missouri, for a patient population diagnosed with non-small cell lung cancer. To evaluate the effectiveness of GPT-4, we compared its results against 2 subject matter experts’ manual annotation. We also conducted a comparative analysis with GPT-3.5-turbo^26^, Flan-T5^27^ (Flan-T5-xl, Flan-T5-xxl), and spaCy (medspaCy, scispaCy), currently frequently used rule-based and machine learning approaches in clinical phenotype extraction. While Flan-T5 models are LLMs, spacy models are two recent and hybrid frameworks that utilize statistical and machine-learning methods in conjunction with rule-based NLP to identify clinical phenotypes. We selected these baseline models based on their inherent capacity for rapid extraction, and their ability to generate results without requiring training or additional fine-tuning.

Our comparison between scispaCy, medspaCy, Flan-T5-xl, Flan-T5-xxl, GPT-3.5-turbo and GPT-4 aims to highlight the strengths and weaknesses of each approach for cancer phenotype extraction from unstructured clinical text, providing valuable insights into their effectiveness and potential use for cases in cancer phenotype extraction from EHR. In evaluating these current approaches for phenotype extraction, we also note their limitations.

## MATERIALS AND METHODS

To extract a detailed representation of specific lung cancer phenotypes, we used GPT-4, available through Microsoft’s Azure OpenAI Service. We compared and evaluated the performance of the current models by comparing true positives (recall) and false positives at the patient-level. The following subsections discuss the datasets, annotation methods, and methodologies used for extracted information, baseline comparison techniques, and evaluation metrics used to quantify differences in results. **Figure 1** illustrates the pipeline we followed for extraction. The study was approved with a waiver of consent by the Washington University in St. Louis Institutional Review Board.

**Figure 1.**
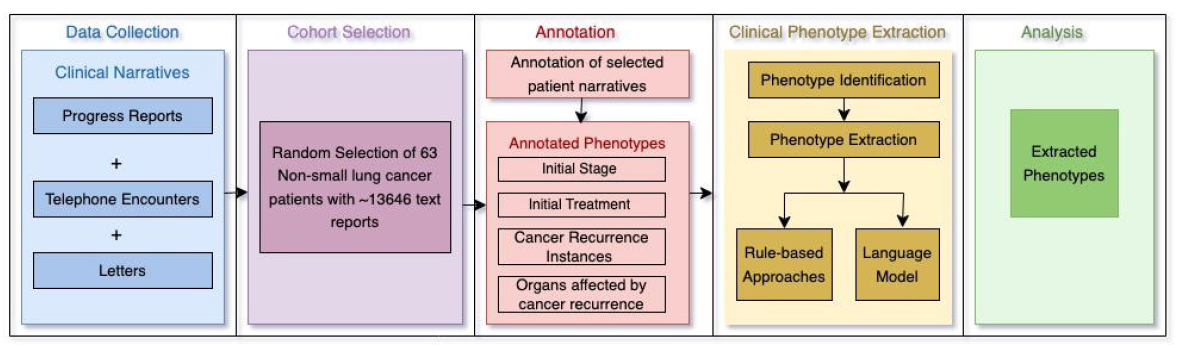
Step-by-step approach to extracting phenotypes. Clinical narratives from the EHR were extracted as part of the data collection process. A subset of the narratives was randomly selected for manual annotation. ScispaCy, medspaCy, GPT-3.5-turbo and GPT-4 models were implemented for phenotype extraction. Extracted phenotypes were compared with the annotations.

### Dataset

Retrospective outpatient and inpatient EHR data were obtained from Washington University Physicians / BJC Healthcare system in St. Louis, Missouri, for all patient encounters with a non-small cell lung cancer diagnosis between 2018-2023. For this study, we extracted a total of 13,646 texts from the EHR of a randomly selected subset of 63 patients.

### Lung cancer phenotypes extraction from the clinical narratives

Our extraction pipeline currently targets four types of phenotypes: cancer stage, cancer treatment (chemotherapy, radiation, surgery), evidence of cancer recurrence, and organs affected by cancer recurrence. We selected these phenotypes based on suggestions from subject matter experts regarding which phenotypes would be most helpful for a proof-of-concept work. The variations extracted for each phenotype are listed in **Table 1**. We attempted to search for all variations of the targeted phenotypes from the corpus.

**Table 1:**
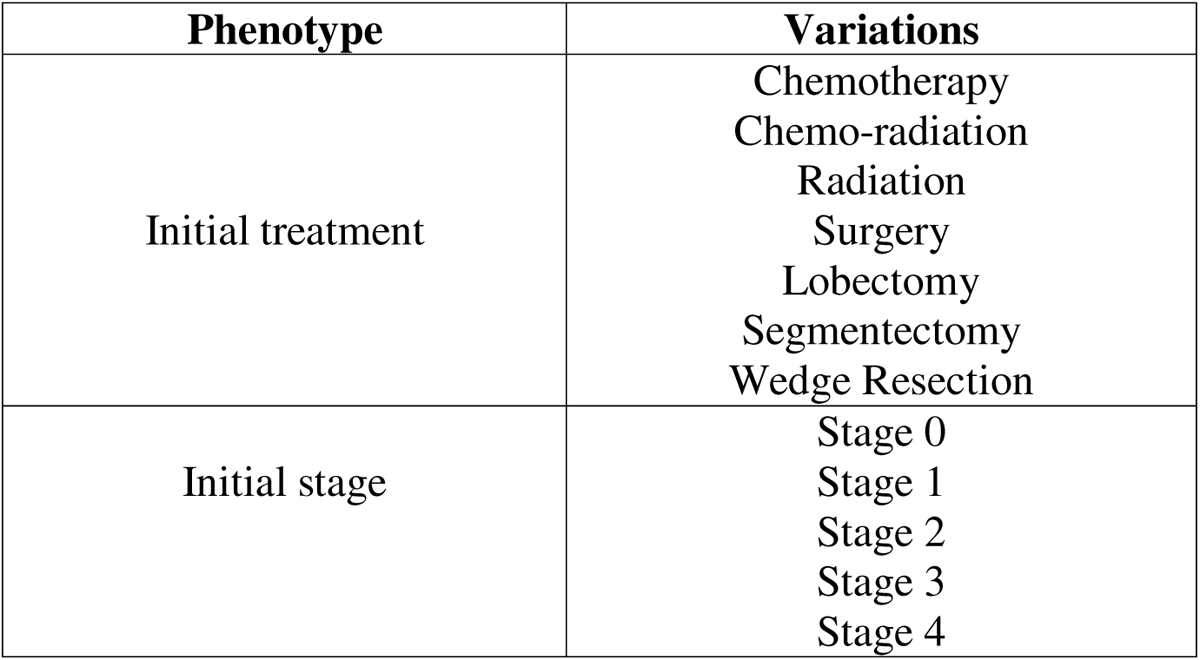
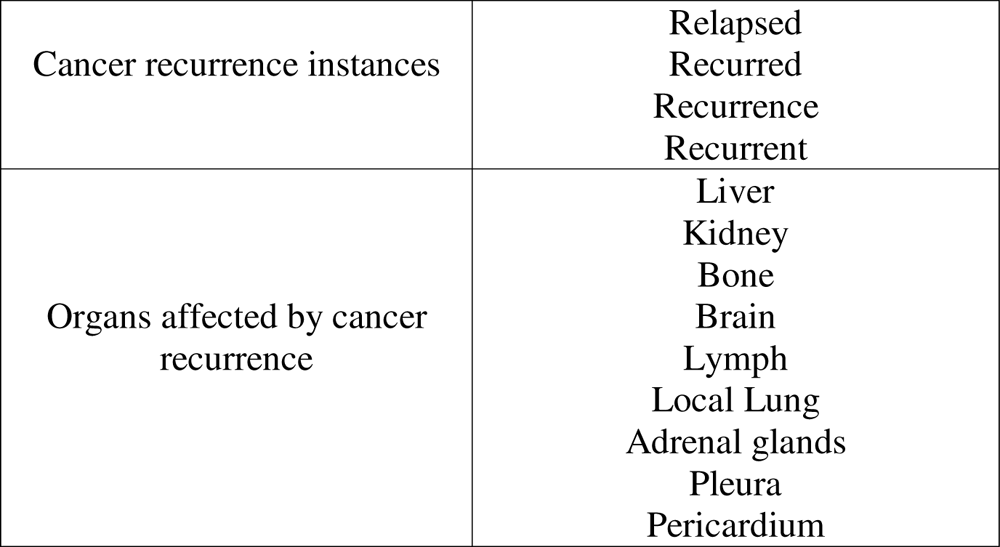
Variations of the relevant phenotypes used in the search for phenotype extraction. All strings were case-insensitive.

### Gold-standard data annotation

The results from the phenotype extraction pipeline for each model were evaluated against gold-standard manual annotation from 2 subject matter experts at the same institution, containing expert determination of initial cancer stage, initial treatment, recurrence instances, and organs affected by cancer recurrence for each patient in the cohort. A Research Electronic Data Capture (REDCap)^28^ form was designed to collect responses from the annotators to comprehensively capture patient phenotypes in a consistent format across annotators. The annotated dataset consisted of the 63 unique patients from the BJC EHR, comprising a total of 13,646 records for all patients.

### Non-small cell lung cancer phenotype extraction and model comparison

We implemented GPT-4 and compared its performance with GPT-3.5-turbo, medspaCy and scispaCy. We constrained ourselves to spaCy models because our initial investigation of two transformer-based language models, T5^29^ and ClinicalBERT^21^, did not effectively capture the necessary phenotypes in its default setting as they are classifier models with labels assigned for each text. We opted against their inclusion in the main manuscript and made comparisons with spaCy’s rule-based and machine learning-based methods which demonstrated better results compared to T5 and ClinicalBERT for baseline comparison. Results utilizing a subset of clinical text for T5 and ClinicalBERT are included in **Supplementary Table 1-2**.

For all models, the input to the models were the phenotypes and their variations. For GPT, we implemented the default zero-shot model where the model input was the text together with the prompt to guide the model for phenotype extraction. We opted for the zero-shot approach to directly compare its performance with the rule-based and machine learning-based approaches. We used the same phenotype variations for extraction across all spaCy model implementations. GPT models did not require inclusion of all phenotype variations. GPT was able to identify stages 0-4 without explicitly mentioning each stage number in the prompt. Similarly, we did not have to explicitly specify each organ type in the prompt to extract organs affected by recurrence.

Our current implementation on GPT and Flan-T5 models focused on capturing both exact and relaxed matches of the phenotype variations mentioned in **Table 1**. Relaxed matches were results that might have some deviations in the results, but they fit the phenotype descriptions keeping the context and the meaning of results the same. Exact matches were exact strings or phrases that were identical between gold standard and LLM.

We performed uncertainty analysis by bootstrapping and calculating confidence intervals to capture model variability and provide insights into the stability of the model’s performance. Bootstrapping resamples model predictions to create a distribution of metrics which can can then be used to estimate confidence intervals. GPT models may exhibit variability in their generated outputs.

### Development of the GPT pipeline as an information extractor to extract each phenotype

GPT-3.5-turbo and GPT-4 are a transformer-based language models developed by OpenAI, trained on large unspecified corpora for multiple NLP tasks. They have been used for natural language generation tasks using their chatCompletion and translations endpoint. Our setup is an adaptation of the sequence labeling task from the chatCompletion framework for phenotype extraction. The sequence labeling setup requires providing context to the model, where the model generates responses that include labeled phenotypes from the clinical notes. The model outputs are the expected phenotypes we are trying to extract. The core idea involves assigning specific labels to individual words or tokens in the clinical notes, capturing the relevant information while retaining the original context. Details on model architecture and training dataset for GPT are provided in **Supplementary Text 1**.

To build the GPT framework, we used Microsoft’s Azure OpenAI Service, which provides REST API access to OpenAI’s language models. We deployed the OpenAI API endpoint via a HIPAA-compliant subscription within Washington University’s Azure tenant. This enabled us to study the performance of GPT on real-world data in a secure and HIPAA-compliant manner. Additionally, we applied for and received an exemption from content filtering, abuse monitoring, and human review of our use of the Azure OpenAI service, which removes the ability of Microsoft employees to perform any form of data review. At the time of our experiments, GPT-3.5-turbo Version 0301 and GPT-4 Version 0613 were the most recent GPT models available.

For phenotype extraction, the model identifies treatment procedures, stage information, and recurrence information from the clinical notes (**Table 1)**. Each note is an input in the prompt along with an instruction to extract the relevant phenotype categories (e.g., treatment, staging) or their sub-categories (e.g., surgery, radiation, chemotherapy, stage numbers) to extract desired information. The primary objective was to compare the performance of GPT-3.5-turbo and GPT-4 in the context of cancer phenotype extraction. Our goal was not to explore different prompting strategies. Therefore, we implemented a zero-shot prompt strategy as our only approach for GPT models. This approach involves providing the model with a single prompt without additional examples or contextual information. The same set of zero-shot prompts was used as input for both GPT-3.5-turbo and GPT-4 to maintain consistency in the evaluation of their performance. We attempted 3-5 variations of prompts for each phenotype, and we selected the prompt that had more accurate results. The final prompts used in this study are reported in **Supplementary Figure 1**. Due to the probabilistic design of GPT models, the output may include extra words or phrases around the actual phenotypes, which were then parsed using regular expressions in the post-processing step (**Supplementary Table 3**). The hyperparameters chosen for the model are reported in **Supplementary Table 4**.

### Development of the spaCy-based NLP pipelines to extract each clinical phenotype using hybrid techniques

In our study, we implemented spaCy’s rule-based and machine learning-based approaches. ScispaCy is a rule-based and Named Entity Recognition (NER)-based Python library for biomedical text processing, which has demonstrated robust results on several Named Entity Recognition (NER) tasks compared to the neural network models of the time.^5^ It is trained on gene data, PubMed articles, medications datasets, and one of their proprietary datasets. We implemented scispaCy version 0.5.2 following the code structure specified in their documentation. For each phenotype of interest, we added specific phenotypes and their corresponding string variations as rules in the pipeline that were then extracted by the model. We incorporated scispaCy’s built-in functions to handle negation and NER. The results were strings extracted from the text and the position of the characters in that text. If a string was not present in the text, the output was null. Finally, the output was mapped into their specific phenotype categories.

MedspaCy is also a rule-based and NER-based Python library that includes UMLS (Unified Medical Language System)^30^ mappings for clinical phenotype extraction. A similar approach was applied for medspaCy (version 1.0.1) as scispaCy. The output from medspaCy was similar to scispaCy, with strings extracted from the text and the position of the characters from that text. The final result from the pipeline were all the strings that medspaCy extracted.

For medspaCy and scispaCy, each existing output string from the clinical notes that matched with phenotype variations was later assigned to the relevant phenotype categories on a patient-level, which were then analyzed as the final extracted phenotypes.

Additional details on the model pipeline and training dataset are provided in **Supplementary Text 2**.

### Development of the Flan-T5 transformer-based model pipeline

Flan-T5 is an open-source language model developed by Google that has been fine-tuned on multiple question answering and text generation tasks. We conducted Flan-T5 experiments with the same prompts that we implemented for the GPT models to make sure the experiment setup was consistent across models. The Flan-T5 models were downloaded from the HuggingFace model hub at https://huggingface.co/google/flan-t5-xxl and https://huggingface.co/google/flan-t5-xl. Similar to the GPT models, we used regular expressions to parse the Flan-T5 output to extract the relevant phenotypes. Additional details on the model architecture and training data are provided in **Supplementary Text 3**.

## RESULTS AND EVALUATION

### Patient Population and Corpus Creation

For the 63 patients selected for this study, average length of each text in corpus is 814 tokens (SD=5022.72). **Table 2** describes the patient demographics used in this study. The unstructured texts for these patients included letters, progress notes, and telephone encounters. Distribution of text types for each phenotype are included in **Supplementary Table 5**. The texts primarily describe patients’ disease trajectory during their visit, ranging from primary cancer diagnosis, cancer stage, treatment type, treatment completion, and cancer recurrence (**Figure 2**).

**Table 2.**
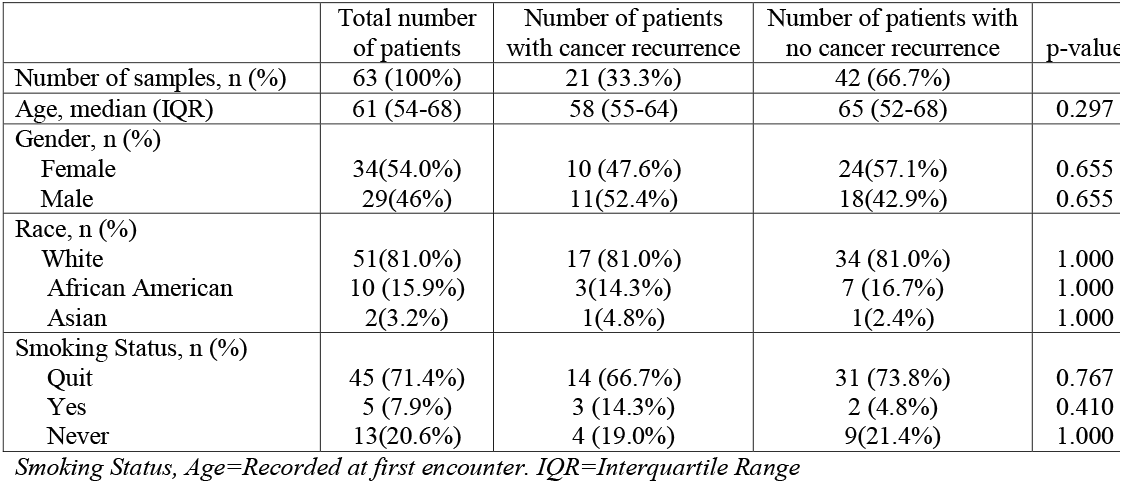
Patient demographics.

**Figure 2.**
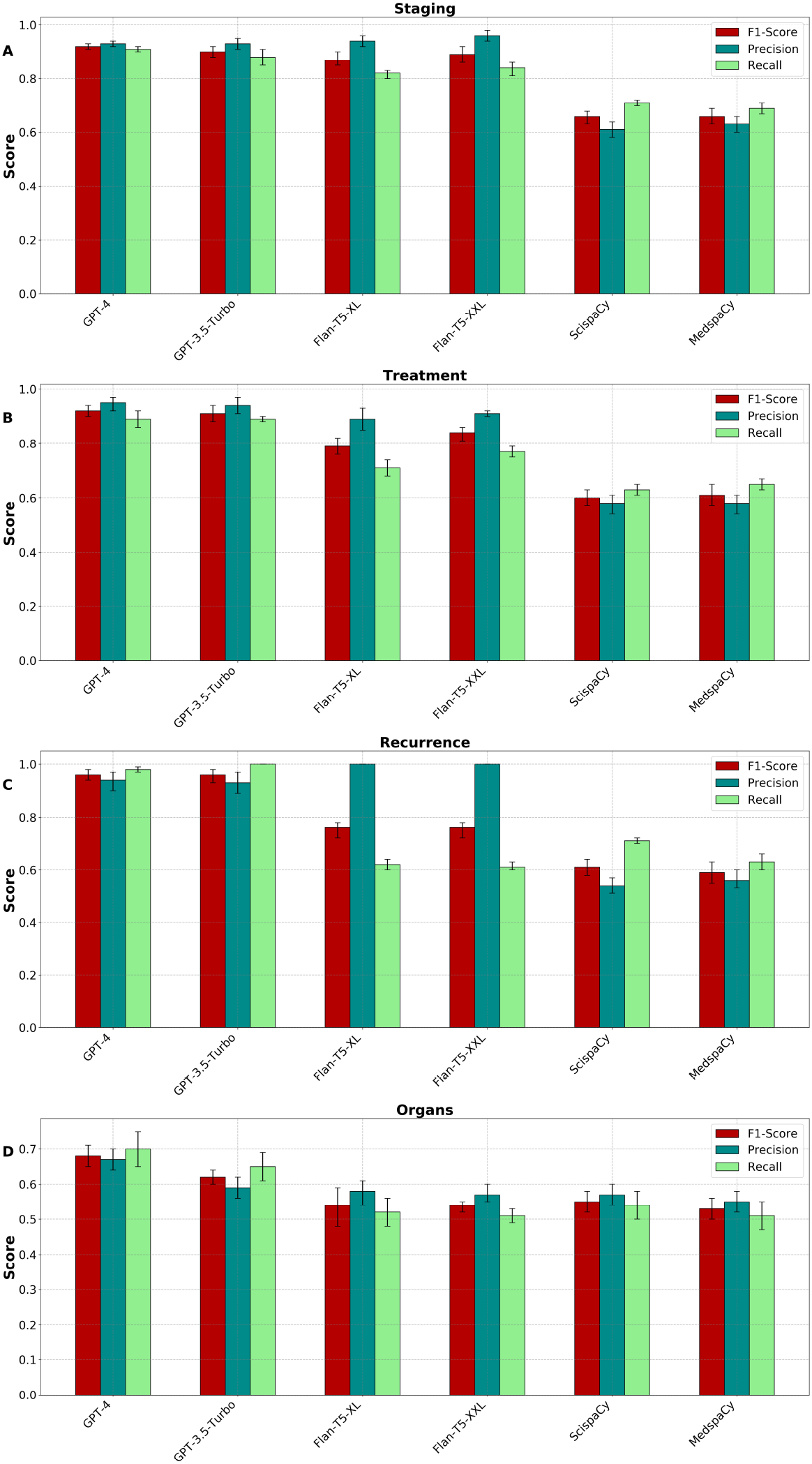
Sample text from unstructured narratives of non-small cell lung cancer patients. The text highlighted in red are the targeted phenotypes for extraction. To protect patient privacy, dates in the figure have been replaced with “XX/XX/XXXX” to protect patient privacy.

### Data Annotation

Inter-annotator agreement initially calculated for each phenotype using Cohen’s Kappa demonstrated high agreement between the annotators (0.68-1.00; **Table 3**). Differences between annotators were resolved through discussions and manual review of the annotations to establish a gold standard for final evaluation.

**Table 3.**
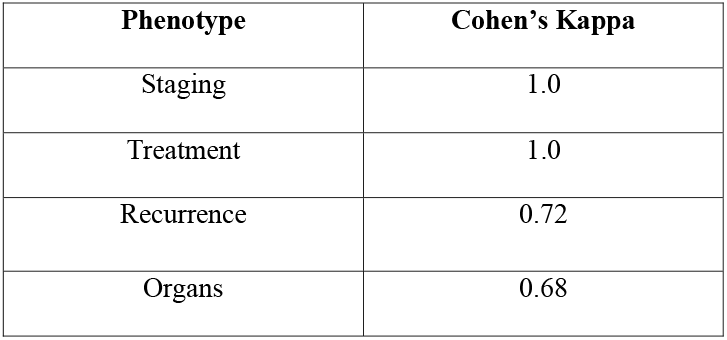
Inter-annotator agreement of manual annotations.

The annotators annotated the narratives by identifying each phenotype from the clinical text for each patient. All the phenotypes mentioned in **Table 1** were identified in the annotator’s annotation, with some phenotypes being identified more frequently than others, depending on the nature of the patient’s disease trajectory. Some patients show cancer recurrence in multiple organs, and the percentage is inclusive of each affected organ. **Table 4** summarizes the frequency of annotations corresponding to each phenotype variation.

**Table 4.**
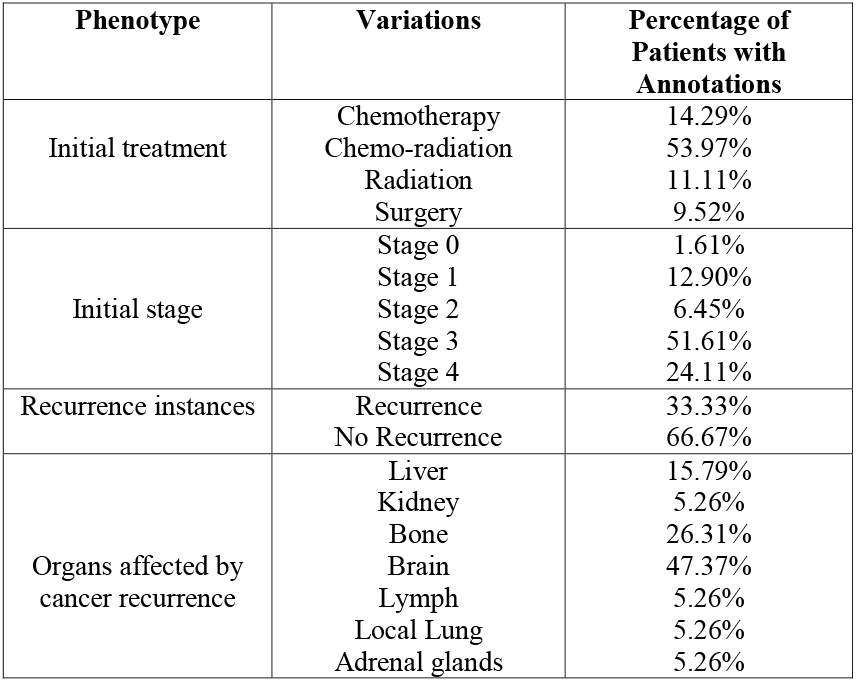
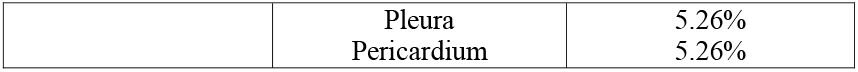
Frequency of annotations corresponding to each phenotype variation identified for each patient within the cohort, based on the available annotations.

We evaluated the performance of each model at identifying the targeted cancer phenotypes (staging, treatment, recurrence, and organs) using precision, recall, and micro-F1 scores to collectively assess the effectiveness of each model in capturing the phenotypes (**Figure 3; Supplementary Table 6**). The inclusion of micro-F1 scores in our evaluation process reflects our emphasis on achieving a balanced assessment, considering both precision (the proportion of correctly identified instances among all instances identified by the model) and recall (the proportion of correctly identified instances among all actual instances) to accurately identify relevant information while minimizing false positives and false negatives, especially in tasks like phenotype extraction from clinical text.

**Figure 3.**
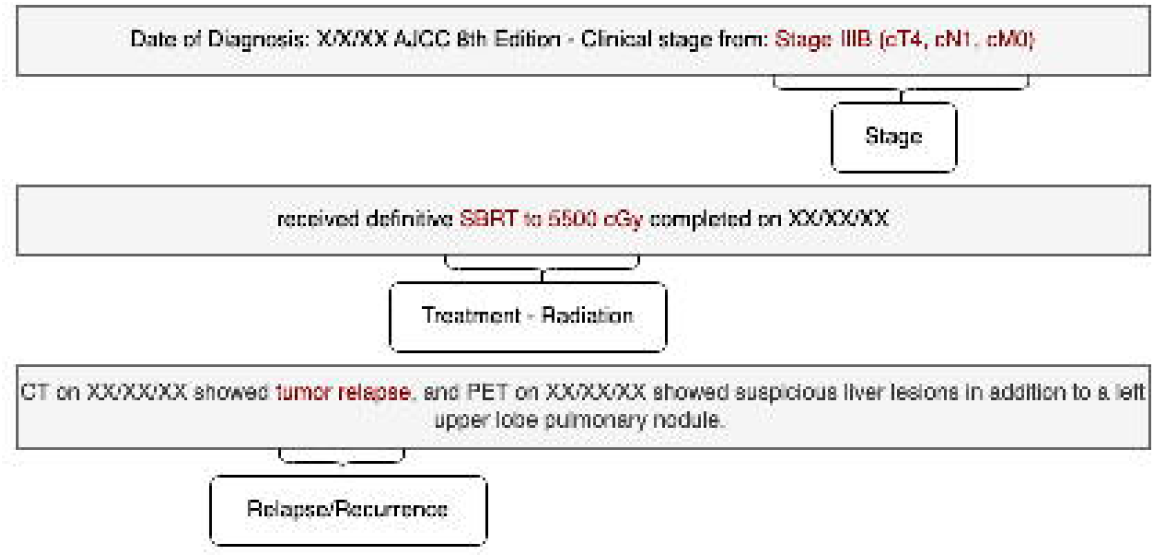
Phenotype extraction performance results for the targeted phenotypes. Comparison of F1-score, precision and recall for scispaCy, medspaCy, Flan-T5-xl, Flan-T5-xxl, GPT-3.5-turbo, GPT-4 models. The figures illustrate the effectiveness of each model in accurately identifying stage (A), treatment (B), recurrence instances (C) and recurrent organs (D) from clinical text data.

### Comparison of Models

The GPT-4 model demonstrated higher F1 scores with high precision and recall, indicating its ability to correctly identify all instances of recurrence, staging, treatment, and organs in the clinical text better than Flan-T5-xl, Flan-T5-xxl, scispaCy and medspaCy. GPT-4 achieved a higher F1 score of 0.96 in identifying recurrence instances compared to staging (0.92), treatment (0.92), and recurrence organs (0.68). GPT-3.5-turbo and GPT-4 had comparable performance across most phenotypes with the recurrence phenotype showing identical F1 score of 0.96. Although scispaCy had lower F1 scores than Flan-T5-xl, Flan-T5-xxl, GPT-3.5-turbo and GPT-4, it outperformed medspaCy in most phenotype extraction tasks. MedspaCy had the lowest F1 score for all phenotypes, suggesting it is less effective at information extraction than other models. This is potentially due to its less advanced NER techniques than scispaCy and GPT models. Evidently, all models were less effective at accurately identifying organ information, likely due to the lack of specific training data for this task. The model-generated output of GPT-3.5-turbo and GPT-4 varied across each run but maintained the underlying meaning of the result across all runs (**Supplementary Table 3**).

### Qualitative Analysis of the Results

We performed a qualitative analysis of the results made by each model in phenotype extraction to better understand their strengths and weaknesses.

GPT-4 was better able to correctly identify cancer phenotypes while minimizing misclassifications, leading to a higher F1 score compared to GPT-3.5-turbo, medspaCy and scispaCy. When comparing GPT-3.5-turbo and GPT-4, we found that both models captured contextual information accurately. However, the generated text from GPT-4 is more relevant to the prompt than the text generated from GPT-3.5-turbo (**Supplementary Table 7**). Upon examining the errors, we observed that GPT models sometimes mislabeled phenotypes when the context was ambiguous, especially when the same sentence discussed multiple phenotypes.

MedspaCy and scispaCy could not identify contextual phenotypes or phenotypes mentioned in a negated context, synonyms not part of the rules, and spelling errors. GPT-3.5-turbo and GPT-4 were far better in these cases. For example, GPT-3.5-turbo and GPT-4 were able to identify “T1c N0 M0” as an indication of a cancer stage, whereas the other models could not identify stage without significant further pipeline engineering. (**Supplementary Table 8**,**9**). This could be due to spaCy’s inability to learn contextual information.

Across all models, the more specifically we defined a phenotype in the prompt or by rules, the better were our chances of identifying it correctly.

## DISCUSSION

Our study highlights GPT-4’s remarkable performance in identifying phenotypes with minimal preprocessing and postprocessing steps compared to rule-based or traditional machine-learning-based algorithms. This aligns well with the established notion that LLMs are data-driven and highly effective even with limited contextual information, unlike rule-based or traditional machine learning algorithms that rely solely on predefined patterns or rules known to researchers or clinicians.^31^

GPT-3.5-turbo performs similarly to GPT-4 for some phenotypes. The choice of GPT-3.5-turbo versus GPT-4 would depend on model run-time and cost of the runs. While GPT-4 is more scalable as its results are more relevant to prompts, GPT-3.5-turbo may be more cost-effective for larger tasks, even when accounting for the additional engineering time necessary to process its output **(Supplementary Table 10)**. Overall, GPT models, with their robust unsupervised pre-training and remarkable pattern recognition capability on tokens, outperform other models as they extract relevant patterns and relationships without being constrained by the need for prior knowledge of explicit patterns, rules, or meaning. Based on the context provided in the prompt, GPT can capture variations in the representation of the clinical phenotypes, making it well suited for information extraction tasks that could extend beyond this study’s focus on its application in oncology.

Our analyses also revealed that GPT demonstrated significantly better performance improvement than the other models, even in its default zero-shot setup without fine-tuning on clinical text. Fine-tuning with clinical text requires additional labeled clinical text, which is not readily available and would have been time-consuming to procure.

For the GPT model outputs, we also obtained varying texts from the API across multiple iterations of the same query despite using the same prompt, suggesting that GPT model might not provide identical results across multiple iterations of the same query. This could be due to its probabilistic design. After analyzing the output texts, we found that all the extracted phenotypes were correctly identified within the text, with only differences in the words and language used.

The comparative analysis also revealed that scispaCy performed better than medspaCy in our study, possibly because of the additional NER components and diverse data sources that it is trained on, in addition to handling the specific type of data that medspaCy is trained on. However, both approaches exhibited limitations in handling complex patterns and context-specific phenotypes. Results from medspaCy and scispaCy also indicate that rule-based models do not handle speculation, and context ambiguity adequately, particularly within complex sentence structures (**Supplementary Table 8-9**).

Furthermore, while medspaCy and scispaCy offer deterministic results based on predefined rules, they fall short of capturing the contextual information required for effective information extraction in clinical text. Because of that, researchers must also have comprehensive knowledge of the phenotypes and variations of the phenotypes for extraction.

Finally, it is worth considering the interpretability aspect of these models. While medspaCy and scispaCy’s rule-based nature allows for more straightforward interpretability, there might be some challenges in interpreting the results of the GPT model due to its unknown internal parameters.

### LIMITATIONS

Despite these promising results, we acknowledge some limitations in this study. We evaluated our results using F1 metrics, which have proven effective in comparing the performance of LLMs to that of rule-based and machine learning-based models for information extraction. However, it is important to reconsider the utility of traditional evaluation metrics when comparing LLM-generated text with human-generated reference text. This is crucial due to the potential discrepancies in reference texts and variations in the representation of results across different LLMs, suggesting that traditional information retrieval metrics may not be well suited for all LLM tasks. Addressing these limitations will be a key focus in our future research.

Additionally, we note that our random selection of a subset of patients may introduce bias and affect model performance. While the dataset was extracted from a 5-year cohort, the evaluation was based on a random subset of patients. Biases in the EHR data and data used for training the models could also lead to limitations in handling diverse clinical text or phenotypes and affecting model performance. Including a larger dataset in future research would address this limitation.

Finally, we acknowledge that our study did not conduct multiple runs of the GPT models due to cost limitations. While some recent work in the non-clinical domain has demonstrated LLMs’ highly consistent results over multiple runs, further research is necessary to determine the optimal number of runs required for reliable clinical phenotype extraction, particularly in the context of lung cancer^32^. Future work will focus on random subsampling on a subset of the data and permutation testing on the subsample to assess model variability.

## CONCLUSION

In conclusion, the study highlights the potential of GPT-4 for accurate phenotype recognition in clinical text. GPT-3.5-turbo model demonstrates performance similar to that of GPT-4. Both GPT models seem to be effective not only for text generation tasks but also surprisingly for information extraction tasks. While medspaCy and scispaCy offer deterministic results and have utility for some tasks, they exhibit limitations in handling complex patterns and context-specific phenotypes. Therefore, leveraging data-driven and contextually aware advanced language models like GPT-3.5-turbo and GPT-4 and addressing their current limitations opens up new possibilities for robust clinical phenotype extraction, ultimately leading to additional insights into patients’ health and improved care.

## ACKNOWLEDGEMENTS

This work was supported by Centene Corporation contract (P19-00559) for the Washington University-Centene ARCH Personalized Medicine Initiative. Jonathan Moran Sierra was supported by NIH/NIGMS T32GM007200.

