## Supplementary File, Supplementary Table, Supplementary Text for "Leveraging GPT-4 for Identifying Cancer Phenotypes in Electronic Health Records: A Performance Comparison between GPT-4, GPT-3.5-turbo, Flan-T5 and spaCy’s Rule-based & Machine Learning-based methods"

**SUPPLEMENTARY MATERIAL**

**Supplementary Table 1.** Sample results from a subset of texts processed using T5. The largest T5 model with 11b parameters was used for this test.

| **Phenotype** | **Prompt+Input Text** | **Output** | **Expected Output** | **Result explanation** |
| --- | --- | --- | --- | --- |
| Stage | **Prompt:** “Extract initial stage information from the following text”  **Section of the input text:** “Diagnosis: Pulmonary adenocarcinoma Stage IV Diagnosed xx/xx/xx PD-L1 = 100% Genomics: FISH: negative for ALK gene rearrangement | 'adenocarcinoma. a a a,: - resection of the cerebellar tumor on xx/xx/xx and -1141.4. No hepatosplenomegaly. No lytic or blastic osseous lesions. No edema. Skin: Denies rash, heartburn, nausea and vomiting. Eyes: Denies blurred vision and' | Stage IV | The model haphazardly summarizes the text without being specific to the context. |
| Treatment | **Prompt: “**Extract initial treatment information from the following text” **Section of the input text:**  “TREATMENT HISTORY: 1. Concurrent chemoradiation with carboplatin and paclitaxel · Thoracic radiation given from 01/15/2020-02/26/2020.” | : Current: CURRENT: /// 2. | Chemoradiation | The model does not generate the expected result |
| Recurrence | **Prompt:** “Identify whether a patient had cancer recurrence/relapse after first treatment completion from the following text. Output: Relapse ‘Yes’ or ‘No’ and relapse/recurrence date” **Section of the input text:** “who developed inflammatory locoregional recurrence with nodal involvement s/p second line chemo (carboplatin/gemcitabine) with minimal pathologic response” | ss// to chemo. -/////e e//oos. | Logoregional recurrence or Yes | The model generated incorrect result when expected to identify recurrence |

**Supplementary Table 2.** Results from a subset of texts processed using clinicalBERT

| **Phenotype** | **Keyword+Input Text** | **Output** | **Expected Output** | **Result explanation** |
| --- | --- | --- | --- | --- |
| Stage | **Keyword:**stage  **Section of the input text:** Staging form: Lung, AJCC 8th Edition - Clinical stage from xx/xx/xx: Stage IIIC | from | Stage III | To search and extract clinicalBERT embeddings that corresponds we set a keyword. Only based on that keyword, ClinicalBERT extracts the desired information using their contextual embeddings. In this text, model extracts the first occurring ‘stage’ and the ‘word’ right after ‘stage’. |
| Stage | **Keyword:** stage  **Section of the input text:**  REASON FOR CONSULTATION: New diagnosis of non-small cell lung cancer. ONCOLOGY HISTORY: Oncology History DIAGNOSIS: cT1N2, Stage IIIA adenocarcinoma of the right upper lobe | IIIA | IIIA | Extracts the expected output |

**Supplementary Text 1. Additional details on the GPT models**

*Model Description*

GPT-3.5 and GPT-4, developed by OpenAI, are transformer-based language model trained for multiple NLP tasks, including natural language generation. Our setup is an adaptation of the sequence labeling task where we provide context as input to the model, and the model generates responses. The context includes the clinical text and prompt where we ask GPT model to extract expected phenotypes from the clinical notes. The models follow transformer model architecture. According to openAI, given both the competitive landscape of developing these models and the safety implications of large-scale models like GPT-4, they are not providing any additional details on the model architecture (including model size), hardware, training compute, dataset construction, and the training method.

*Training Dataset*

The models were trained using publicly available data (such as internet data) as well undisclosed licensed data. According to OpenAI, it is a web-scale corpus of data including correct and incorrect solutions to math problems, weak and strong reasoning statements, self-contradictory and consistent statements, and statements representing varying ideologies. The data used to train the model is from until September 2021. Data generated after September 2021 was not used to train the model.

*Framework Setup*

To build the GPT framework, we used Microsoft’s Azure OpenAI Service, which provides REST API access to OpenAI’s language models. We deployed the OpenAI API endpoint into a HIPAA-compliant subscription within Washington University’s Azure tenant. This enabled us to study the performance of GPT in a secure and HIPAA-compliant manner. Additionally, we applied for and received an exemption from content filtering, abuse monitoring, and human review of our use of the Azure OpenAI service, which removes the ability of Microsoft employees to perform any form of data review. At the time of our experiments, GPT-3.5 Version 0301 and GPT-4 Version 0613 were the most recent GPT models available. The openAI version 0.28.1 was used for the experiments in this study.

**Supplementary Figure 1.** Prompts selected in the GPT Model to extract all phenotypes.

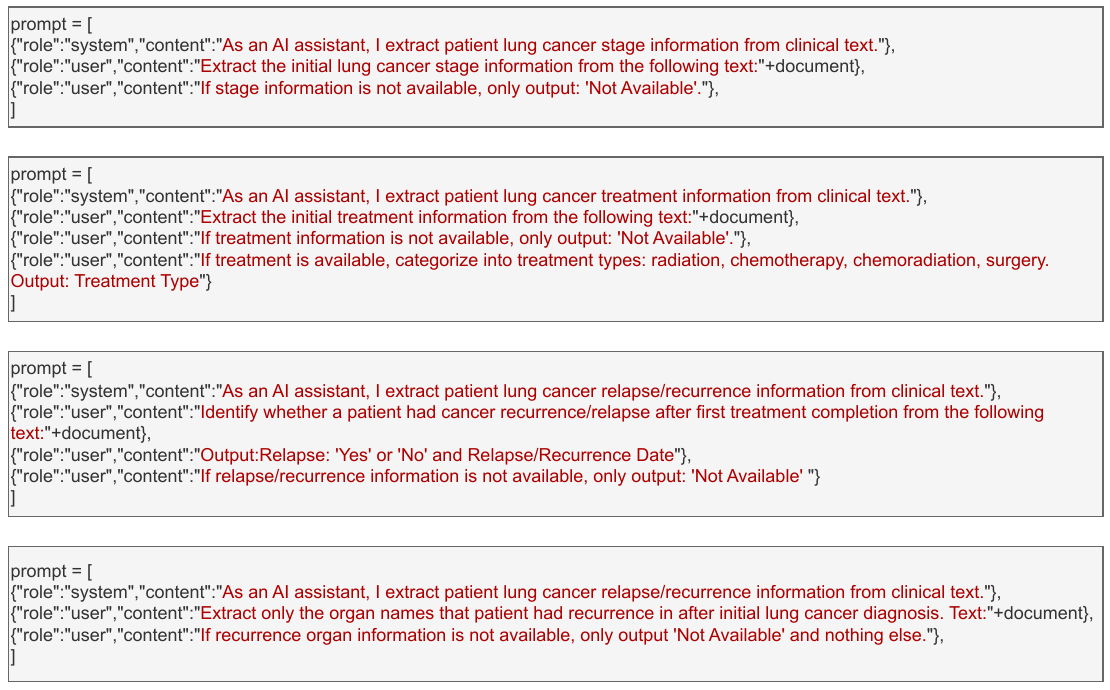

**Supplementary Table 3.** GPT Output Variations

| **Phenotype** | **Output Variations** |
| --- | --- |
| Staging | Output: Stage IIIA,  Output: T1 N2 M0,  Output: Stage 4,  The initial lung cancer stage mentioned in the text is Stage IIIB,  Output: Stage T4N3M0 (IIIC) |
| Treatment | Treatment: Radiation therapy,  Treatment: Chemoimmunotherapy (carbo/pem/pembro),  Treatment: Carboplatin plus pemetrexed and pembrolizumab,  Treatment: PACLItaxel / CARBOplatin with Concurrent Radiation,  Treatment: Taxol/Carbo,  Treatment: Concurrent chemoradiation, 60 Gy in 30 fractions (CoT xx/xx/xx) with weekly carbo/taxol |
| Cancer recurrence | Relapse/recurrence information: Disease progression was noted in mediastinal nodes and the calvarium on CT imaging on xx/xx/xx,  Relapse/recurrence information: Locoregional recurrence of non-small cell lung cancer (NSCLC) with possible NE differentiation,  Relapse/recurrence information: Locally advanced non-small cell lung cancer with local regional recurrence |
| Recurrence organs | Relapse/recurrence organ: Brain, bone, and lung,  Relapse/recurrence organ: Mediastinal nodes and the calvarium,  Relapse/recurrence organ: Left supraclavicular lymph nodes,  Relapse/recurrence organ: The relapse/recurrence is in the right lower lobe of the lung,  Right upper lobe lung mass, hepatic segment 4 |

**Supplementary Table 4.** Hyperparameters used in the model.

| **Hyperparameter** | **Value** |
| --- | --- |
| Tokenization and Context Window | 200 tokens |
| Temperature (Randomness of the model output) | 0 |
| Top p (Top-K Sampling Technique) | 0.95 |
| Presence_penalty (Penalty to discourage model from generating responses that contain certain specified tokens) | -1.0 |

**Supplementary Text 2. Additional details on the spaCy models**

**ScispaCy**

*Model Description*
 ScispaCy is a specialized NLP library for processing biomedical texts which builds on the spaCy library. It was first published in 2019. The model en_core_sci_md within scispaCy have a larger vocabulary and include word vectors compared the original spaCy library which is trained for POS tagging, dependency parsing, and Named Entity Recognition using datasets relevant to biomedical text. The tokenization module within scispaCy has also been improved with additional rules. The spaCy was designed for general NLP tasks covering multiple domains. ScispaCy models with its added features improves on the accuracy and terminology recognition for healthcare text. At the time of our experiments, version 0.5.3 was the most recent model for use.

*Framework Setup*

We input the text into the scispacy model where individual components of the model was added in to process, tokenize, tag, parse, and normalize the input to perform scientific NER and extract the phenotypes as the input.

*Training Dataset*
 ScispaCy has incorporated training data from the OntoNotes 5.0 corpus when training the dependency parser and POS tagger. To train the dependency parser and part of speech tagger in both released models, they convert the treebank of McClosky and Charniak, 4 which is based on the GENIA 1.0 corpus to Universal Dependencies v1.0 using the Stanford Dependency Converter. As this dataset has POS tags annotated, we use it to train the POS tagger jointly with the dependency parser in both released models. Finally, they also leveragedPubMed abstracts for training the models. In addition, they also include relevant named entities linked to their Medical Subject Headings (MeSH terms) as well as chemicals and drugs linked to a variety of ontologies, as well as author metadata, publication dates, citation statistics and journal metadata.

To increase the robustness of the dependency parser and POS tagger to generic text, we make use of the OntoNotes 5.0 corpus when training the dependency parser and part of speech tagger The OntoNotes corpus consists of multiple genres of text, annotated with syntactic and semantic information, but we only use POS and dependency parsing annotations in this work. The main NER model in both released packages in scispaCy is trained on the mention spans in the MedMentions dataset. They also have finer-grained NER techniques embedded trained on BC5CDR (for chemicals and diseases), CRAFT (for cell types, chemicals, proteins, genes; Bada), JNLPBA (for cell lines, cell types, DNAs, RNAs, proteins) and BioNLP13CG (for cancer genetics), respectively.

**MedspaCy**

*Model Description*
 The default spaCy tokenizer is not trained on clinical text. The major drawback of the default spaCy tokenizer for clinical text processing is that it is not trained on clinical text. Additionally, it has a variety of rules designed to handle text sourced online, including many rules that mitigate excess tokenization of URLs. These rules prevent splitting sequences of alphanumeric characters and punctuation into multiple tokens. However, URLs are relatively uncommon in clinical text but typos and using punctuation to delineate document structures are common. The tokenizer included in medspaCy implemented custom rules to handle punctuation and inconsistent use of whitespace that are common in clinical notes. It was published in 2021.

Added features in medspacy were Tokenization, sentence split, Sentence Detection, NER techniques, Section Detection, Concept extraction, UMLS mapping, Contextual analysis, pre-post processing utilities. So, there are overlapping and non-overlapping components between scispacy and medspacy. Medspacy which came after scispacy does not have baseline comparison results to each other.

*Training Data*
 From the documentation, it seems like medpsacy’s implementation was on top of spacy’s with added features. With that, the assumption is that there was no additional clinical data added to the actual model. However, for every added feature like section detection, concept extraction, medspacy utilizes LOINC, and a sample of UMLS ontologies to map concepts and improve the extraction results.

**Supplementary Text 3. Additional details on the Flan-T5 models**
*Model Description*
 Flan-T5 is an open-source language model built upon the T5 encoder-decoder archictecture and developed by Google. It leverages instruction fine-tuning on several tasks. The model was built by extending instruction finetuning by scaling the number of finetuning tasks, scaling the size of the model, and finetuning on chain-of-thought (COT) datasets. The performance evaluation also indicates enhanced generalizability across different downstream tasks, including a range of few-shot, zero-shot, and CoT tasks.

*Training Data*

The model is trained on a large dataset that includes data from various natural language processing tasks and coding problems. The data includes more than 1800 inference and training tasks dataset. The model also utilizes 9 chain-of-thought(COT) datasets, which includes answers and reasoning for answers from annotators. The Flan-T5 is trained on instructions referencing these COT annotations, allowing to potentially learn and apply reasoning skills to unseen tasks.

**Supplementary Table 5. Distribution of the text types from where phenotypes were extracted.**

| **Phenotype** | **Text Type** | **Count** |
| --- | --- | --- |
| Treatment | Letter | 22 |
|  | Progress Notes | 40 |
|  | Telephone Encounter | 1 |
| Staging | Letter | 33 |
|  | Progress Notes | 17 |
|  | Telephone Encounter | 2 |
| Relapse Instances | Letter | 13 |
|  | Progress Notes | 8 |
| Recurrence Organs | Letter | 13 |
|  | Progress Notes | 8 |

**Supplement Table 6**: Phenotype extraction performance results for all models. CI=Confidence Interval

| **Approach** | **Phenotype** | **F1-Score  (Point Estimate, 95% CI)** | **Precision (Point Estimate, 95% CI)** | **Recall (Point Estimate, 95% CI)** |
| --- | --- | --- | --- | --- |
| **GPT-4** | Staging  Treatment  Recurrence  Organs | 0.92 (0.91, 0.93)  0.92 (0.90, 0.94)  0.96(0.94,0.98)  0.68(0.65,0.71) | 0.93(0.92, 0.94)  0.95(0.92, 0.97)  0.94(0.90, 0.97)  0.67(0.64, 0.70) | 0.91(0.90, 0.92)  0.89(0.86, 0.92)  0.98(0.97, 0.99)  0.70(0.65,0.75) |
| **GPT-3.5-turbo** | Staging  Treatment  Recurrence  Organs | 0.90(0.88,0.92)  0.91(0.88, 0.94)  0.96(0.93,0.98)  0.62(0.60,0.64) | 0.93(0.91, 0.95)  0.94(0.91, 0.97)  0.93(0.89, 0.97)  0.59(0.56, 0.62) | 0.88 (0.85,0.91)  0.89(0.88,0.90)  1.00(1.00,1.00)  0.65(0.61,0.69) |
| **Flan-T5-xl** | Staging  Treatment  Recurrence  Organs | 0.87 (0.85,0.90) 0.79 (0.76,0.82)  0.76 (0.72, 0.78)  0.54 (0.48, 0.59) | 0.94 (0.92,0.96) 0.89 (0.85,0.93)  1.00(1.00,1.00)  0.58(0.54,0.61) | 0.82 (0.80,0.83)  0.71(0.68,0.74)  0.62(0.60,0.64)  0.52 (0.48,0.56) |
| **Flan-T5-xxl** | Staging  Treatment  Recurrence  Organs | 0.89(0.86,0.92)  0.84 (0.81,0.86)  0.76 (0.72,0.78)  0.54(0.52,0.55) | 0.96(0.94, 0.98)  0.91 (0.90,0.92)  1.00 (1.00,1.00)  0.57 (0.55,0.60) | 0.84(0.81,0.86)  0.77(0.75,0.79)  0.61(0.60,0.63)  0.51(0.49,0.53) |
| **scispaCy** | Staging  Treatment  Recurrence  Organs | 0.66(0.63,0.68)  0.60(0.57, 0.63)  0.61(0.58, 0.64)  0.55(0.52, 0.58) | 0.61(0.58, 0.64)  0.58(0.54, 0.61)  0.54(0.51, 0.57)  0.57(0.54, 0.60) | 0.71(0.70,0.72)  0.63(0.61,0.65)  0.71(0.70,0.72)  0.54(0.50,0.58) |
| **medspaCy** | Staging  Treatment  Recurrence Organs | 0.66(0.63, 0.69)  0.61(0.57, 0.65)  0.59(0.55, 0.63)  0.53(0.50, 0.56) | 0.63(0.60, 0.66)  0.58(0.54, 0.61)  0.56(0.53, 0.60)  0.55(0.52, 0.58) | 0.69(0.67,0.71)  0.65(0.63, 0.67)  0.63(0.60,0.66)  0.51(0.47,0.55) |

**Supplementary Table 7.** Manual review of the true positive results from a subset of texts processed using GPT-3.5-turbo and GPT-4

| **Model** | **Phenotype** | **Input Text** | **Output** |
| --- | --- | --- | --- |
| GPT-3.5 -turbo | Stage | **Snippet of the input text:** The immunoprofile is consistent with an adenocarcinoma (Missouri Baptist Hospital, MC18-1670). T1N2M0 at diagnosis Genomic and molecular markers | The initial lung cancer stage mentioned in the given text is T1N2M0. |
| GPT-4 | Stage | **Snippet of the input text:** The immunoprofile is consistent with an adenocarcinoma (Missouri Baptist Hospital, MC18-1670). T1N2M0 at diagnosis Genomic and molecular markers | T1N2M0 |
| GPT-3.5-turbo | Stage | **Snippet of the input text**: Referral done and faxed | I'm sorry, but it is not possible to extract lung cancer stage information from the given text "Referral done and faxed". This text does not contain any information related to the patient's lung cancer stage. Therefore, the output will be "Not Available". |
| GPT-4 | Stage | **Snippet of the input text:** Referral done and faxed | Not Available |
| GPT-3.5-turbo | Treatment | **Mention of treatments in the input Text:** Concurrent radiation with carboplatin and paclitaxel A. Week one: | Model Output: Radiation  Expected output: Chemo-radiation |
| GPT-4 | Treatment | **Mention of treatments in the input Text:** T3 Nx M0 primary squamous cell carcinoma of the right upper lobe of the lung, with right suprahilar involvement TREATMENT HISTORY: 1. Concurrent chemoradiation with carboplatin and paclitaxel | Model Output: Concurrent chemoradiation with carboplatin and paclitaxel |
| GPT-4 | Recurrence | **Mention of recurrence in the input text:** Radiology Progress Note Interval History: 67 year old man with laryngeal cancer recurrence with reduced | Model Output: No. The patient had laryngeal cancer recurrence. |

**Supplementary Table 8.** Qualitative results from an instance of a patient visit for scispaCy

| **Phenotype** | **Mention of stage and treatments in the input text** | **Output** | **Expected Output** | **Error explanation** |
| --- | --- | --- | --- | --- |
| Staging | Assessment: xx xx is a xx y.o. female with left upper lobe adenocarcinoma, clinical stage T1c N0 M0. | Null | T1c N0 M0 | The model made an error because it was not able to correctly identify the TNM staging of the patients which are documented in different forms (example: T1c N0 M0, T1N0M0, cT2bN2M0, T1b N0 M0) |
| Treatment | The CT shows that a lobectomy would not be sufficient- she would need more than a lobectomy with some partial or complete removal of part of the LLL to get a complete resection. Summary: Stage IIIa (T2N2) with some DOE and some objective evidence of reduced lung capacity. I think the best approach is non-surgical therapy with combined chemo and radiation. | lobectomy | Chemo-radiation | The surgical treatment was a possible treatment, and not a definite one which the model was not able to capture. |
| Recurrence | If there is evidence of disease recurrence, then would re-induce remission with steroids | recurrence | Null | The recurrence instance was a possibility. There was no certainty about the patient having recurrence which the model was not able to identify |
| Organs | Diabetes Mother • Brain Aneurysm Sister passed away | Brain | Null | Recurrence did not occur in the brain for the patient. In this context, there was a family history of brain aneurysm |

**Supplementary Table 9.** Qualitative error analysis on a selected sample of sentences from an instance of a patient visit for medspaCy

| **Phenotype** | **Mention of recurrence in the input text:** | **Output** | **Expected Output** | **Error explanation** |
| --- | --- | --- | --- | --- |
| Recurrence | She has not had a recurrence of chest tightness. | Recurrence | Null | The model made an error by incorrectly extracting a negated instance of chest tightness for instances when it was expected not to extract any phenotype, as the input text does not indicate cancer recurrence. |
| Organs | Please specify the stage of Chronic Kidney Disease and document in the medical record and on the form below. ___ Chronic Kidney Disease, Stage II (Mild) | Kidney | Null | The model identified kidney as one of the organs for recurrence, when in fact it was a diagnosis for chronic kidney disease. Because of the rule-based setup, medspaCy is unable to identify contextual information |
| Organs | In addition to his vascular disease, the radiologist also noted some fatty changes in his liver. | Liver | Null | The model extracted liver as the organ for recurrence. However, the sentence does not indicate cancer recurrence in the liver |
| Treatment | T3 Nx M0 primary squamous cell carcinoma of the right upper lobe of the lung, with right suprahilar involvement TREATMENT HISTORY: 1. Concurrent chemoradiation with carboplatin and paclitaxel | chemoradiation | chemoradiation | The model was able to identify the treatment as it was explicitly mentioned in the text |
| Staging | Assessment: xx xx is a xx y.o. female with left upper lobe adenocarcinoma, clinical stage T1c N0 M0. | Null | T1c N0 M0 | The model made an error because it was not able to correctly identify the TNM staging of the patients which are documented in different forms (example: T1c N0 M0, T1N0M0, cT2bN2M0, T1b N0 M0) |
| Staging | AJCC 8th Edition - Clinical: FIGO Stage IVB | Stage IVB | Stage IVB | Model is able to identify explicit mentions of the stage which was specified in the rules |

**Supplement Table 10.** Time and cost to run the phenotypes with GPT models

|  | **Runtime** | |
| --- | --- | --- |
| **Models** | **Time (Average in hours)/ phenotype** | **Total Time (in hours)**  **(Phenotypes=4)** |
| GPT-3.5-turbo | 5.92 | 23.68 |
| GPT-4 | 8.54 | 34.16 |

|  | Rate | Token Length | Cost (in Dollars)/ phenotype | Total Cost  (Phenotypes = 4) |
| --- | --- | --- | --- | --- |
| GPT-4 | Input ($0.03/1k tokens) | 11092178 | 332.76 | 1331 |
|  | Output ($0.06/1k tokens) | 1364600 | 81.876 | 327.504 |
|  | Total | 12456778 | 414.636 | 1658.504 |
| GPT-3.5-turbo | Input ($0.003/1k tokens) | 11092178 | 33.27 | 133.11 |
|  | Output ($0.004/1k tokens) | 1364600 | 5.46 | 21.84 |
|  | Total | 12456778 | 38.73 | 154.95 |
